# Engineering the Compression of Sequencing Reads

**DOI:** 10.1101/2020.05.01.071720

**Authors:** Tomasz Kowalski, Szymon Grabowski

## Abstract

**Motivation:** FASTQ remains among the widely used formats for high-throughput sequencing data. Despite advances in specialized FASTQ compressors, they are still imperfect in terms of practical performance tradeoffs.

**Results:** We present a multi-threaded version of Pseudogenome-based Read Compressor (PgRC), an in-memory algorithm for compressing the DNA stream, based on the idea of building an approximation of the shortest common superstring over high-quality reads. The current version, v1.2, practically preserves the compression ratio and decompression speed of the previous one, reducing the compression time by a factor of about 4–5 on a 6-core/12-thread machine.

**Availability:** PgRC 1.2 can be downloaded from https://github.com/kowallus/PgRC.

## 1 Introduction

Raw sequencing data in FASTQ format still contribute significantly to the vast volumes of genomic resources. A number of dedicated compression solutions have been proposed in the last decade, mostly focusing on the short reads (typically of length 100 or 150 bp). Most recent algorithms allow removing the redundancy in overlapping reads, to name SCALCE (Hach *et al*., 2012), ORCOM (Grabowski *et al*., 2015), FaStore (Roguski *et al*., 2018), SPRING (Chandak *et al*., 2018b), Minicom (Liu *et al*., 2018), FQSqueezer (Deorowicz, 2020) and PgRC (Kowalski and Grabowski, 2020). In our opinion, FaStore and SPRING are reasonable practical choices, the former being faster and the latter usually offering higher compression ratios, but if we value compression efficiency even more, the choice may be between PgRC and FQSqueezer. PgRC wins in the compression ratio by up to 15% over SPRING and is rather fast in the decompression, although not in the compression, being a single-threaded program. On the other hand, FQSqueezer is even stronger in compression (by up to around 10%), yet its computational requirements are significant, being many times slower in decompression than PgRC, due to the symmetric nature of the underlying PPM compression scheme. More details about these (and other) tools can be found, e.g., in (Kowalski and Grabowski, 2020).

In this work, we present an improved version of PgRC (v1.2), where the main changes are focused on speeding up the compression. One is parallelization of almost all of its phases, another is a simple preprocessing technique allowing to find LZ-matches much faster. Please note that PgRC is (still) not a full-fledged FASTQ compressor, as it only compresses the DNA stream. Still, we believe that in the current release it may be a relevant yardstick for measuring the performance of future DNA read compressors.

## 2 Materials and Methods

PgRC works in several stages (we use this word interchangeably with “phases” throughout the paper), which are now briefly presented; more details can be found in (Kowalski and Grabowski, 2020, Sect. 2). The parallelization, added for most stages, is obtained with aid of the OpenMP library, with the exception of the backend compressor (LMZA), which programmatically supports two threads. From the input FASTQ its DNA stream is extracted (in a single-threaded phase), and an approximation of the shortest common superstring (called later a “pseudogenome”, or pg in short) over the high-quality reads built. The high-quality reads are found during an early step of the pg construction, which is later called “read set division”. They are those reads which produce long enough both left and right overlaps with regard to other reads (please note that we do not use quality scores in distinguishing between high- and low-quality reads). Once the high-quality reads are separated from the low-quality ones (and also from the reads containing the N symbols), the pseudogenome over them is constructed, denoted as *P G*_*hq*_. Finding read overlaps in a multi-threaded implementation required care, since a naive parallel matching may lead to cycles among overlapping reads. In the next stage, the reads which do not participate in *P G*_*hq*_ (and thus generally being of lower quality) are mapped onto *P G*_*hq*_ with several allowed mismatches (i.e., Hamming errors). More concretely, we allow up to *lread*_*len/MJ* mismatches per aligned read, where *M* is 5 by default. This means that for reads of length 100 we allow up to 20 mismatches between the read and the area of *P G*_*hq*_ it is aligned to.

Each successfully aligned read is represented with several items of data, sent to multiple (conceptual) streams and later encoded and compressed using custom techniques. Those data include the offset (position) in *P G*_*hq*_ of the aligned read, its number of mismatches, their positions (encoded differentially), the read’s symbols at the mismatching positions, and a flag to tell if the read is mapped forwardly or reverse-complemented. The read alignment stage is now multi-threaded.

From the low-quality reads which have not mapped onto *P G*_*hq*_, we create two pseudogenomes, *P G*_*lq*_ and *P G*_*N*_, where the latter is based on reads containing symbols N. The construction of those pseudogenomes is identical as for *P G*_*hq*_ and also multi-threaded.

The high-quality pseudogenome, *P G*_*hq*_, contains specific redundancy that wouldn’t be removed in the final stage when a general-purpose compression algorithm is applied. We mean here reverse-complemented matches of relatively long strings. We find those matches quite efficiently using a hash table and a sparse sampling technique (Grabowski and Bieniecki, 2019). The hash table is built using multiple threads. The implementation of the match search itself is currently serial, but we do not anticipate problems with parallelization (although the expected overall speedup will be rather small). The mentioned sparse sampling technique is used twice (although with different parameters), in the read alignment and the reverse-complemented matching phase, but no collisions during the hash table construction are handled across different threads. This means that some of the matches which would be found using a single-threaded implementation may now be missed, but in practice it is a rare situation, and our design choice speeds things up.

What is left is backend compression on the resulting components. Like in the previous PgRC version, we stick to LZMA for the pseudogenomes (which constitute the lion’s share of PgRC’s by-products) and PPMd for some of the remaining components (the other ones are LZMA-compressed), from the widely used 7zip compression tool. There are two changes, though. First (quite trivial) is that we run the LZMA algorithm with two threads, as supported by the algorithm’s API. The other change is intended to reduce the LZMA compression time via using a more dense (packed) input representation of the pseudogenomes. Following the idea from (Bonfield and Mahoney, 2013, p. 8), we pack together a variable number (up to 4) of successive DNA bases into a byte, in a way that facilitates self-synchronization, which is important not to lose (or shorten by much) too many matches in the further, compression stage (Fig. 1 illustrates). Below we present some details about our packing scheme.

**Fig. 1.**
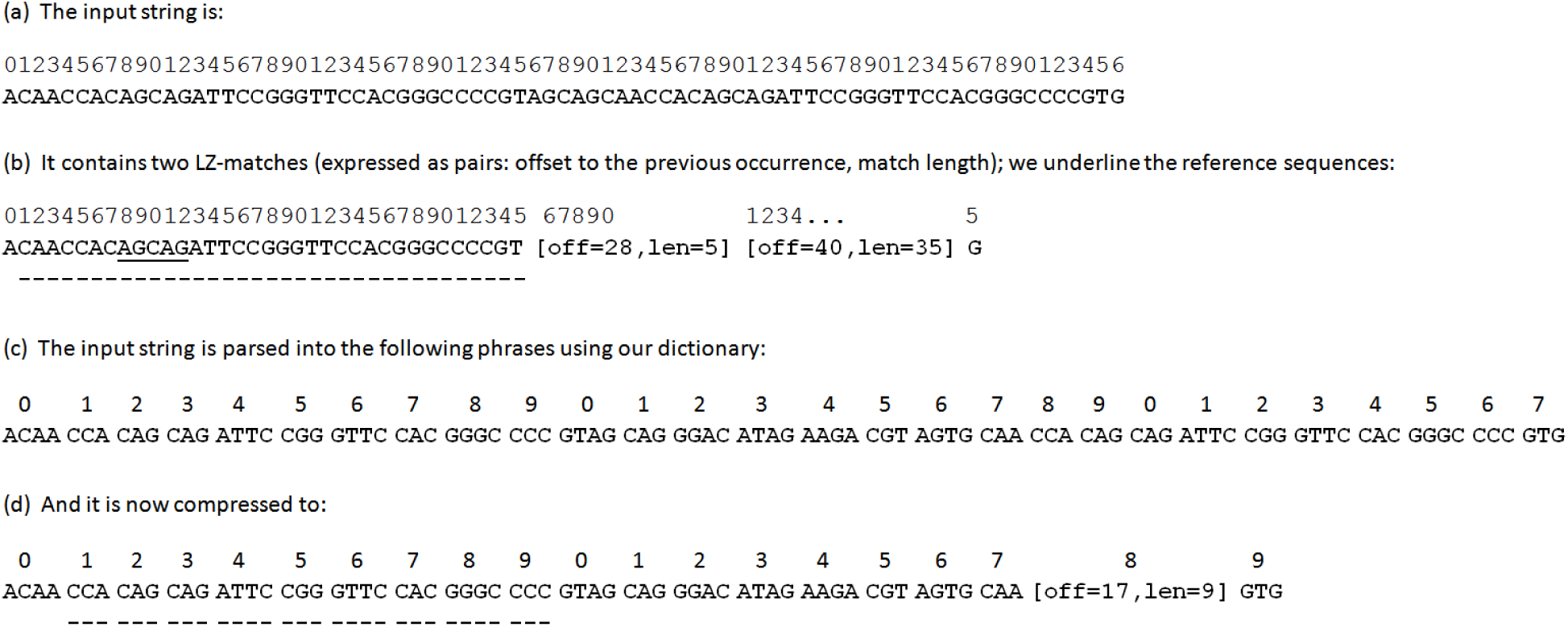
An example of LZ-matches in the natural representation of the DNA string (figure (a) and (b)), and the impact of variable-length packing (figure (c)) on further LZ-matching (figure (d)). The minimum LZ-match length in the original representation is assumed to be 5. Instead of two LZ-matches, of length 5 and 35 symbols, in the compact representation we have only one match spanning over 30 original symbols.

The dictionary of 242 phrases (of length from 1 to 4 bases) is arranged according to their reversals (e.g., phrase TAA is just before phrase ATAA), with some locations in the range 0 … 255 unused. Due to some technical reasons related to the backend LZMA compressor (and thus improving the compression a bit), we start each group of phrases ending with the new standard symbol (A, C, G, T) at a multiple of 64. Some phrases include the symbols N and %, the latter signaling reverse-complemented matches (see the constant string PgSAHelpers::VarLenDNACoder::AG_EXTENDED_CODES in the sources for details). The encoder works greedily, always finding the longest prefix of the remaining text corresponding to a phrase in the dictionary. Both the text encoding and decoding using the presented variable-length scheme are rather fast (achieving, roughly speaking, around 500 MB/s on our machine) and in the current implementation are single-threaded.

Transforming the DNA sequence to such packed representation usually has a minor (positive or negative) impact on the final compression, although on one of our datasets the loss is quite significant. Overall, however, we found using it a viable time-compression tradeoff.

## 3 Results

PgRC was tested on a Linux machine equipped with an 6-core Intel Core i7-7800X 3.5 GHz CPU, 128 GB of DDR4-RAM (CL 16, clocked at 2666 MHz) and a 5 TB HDD (Toshiba X300 HDWE150EZSTA).

The test collection consists of a number of real genome and transcriptome sequencing datasets, used for benchmarking in prior works on FASTQ compression. Their basic characteristics are given in Table 1. All, original and compressed, dataset sizes are given in GBytes (or Gigabases), where *G* = 10^9^. SPRING (version from 2019-Sep-12, https://github.com/shubhamchandak94/Spring) and PgRC 1.2 compressors were run with 12 threads (the test CPU sports 12 hardware threads, thanks to hyper-threading) and other settings default. PgRC 1.1 is a single-threaded implementation.

**Table 1.** Datasets used in the experiments

| SE_dataset | gen. length<br>(Mbases) | reads<br>(M) | coverage<br>(times) | read<br>length | PE_dataset | gen. length<br>(Mbases) | read pairs<br>(M) | coverage<br>(times) | read<br>length |
| --- | --- | --- | --- | --- | --- | --- | --- | --- | --- |
| SRR870667_1 (T. cacao) | ~340 | 69.2 | 22 | 108 | ERP001775 (H. sapiens) | 3,234 | 439 | 27 | 101 |
| SRR445724 (H. sapiens, RNA-Seq) | — | 50.9 | — | 100 | ERR174310 (H. sapiens) | 3,234 | 208 | 13 | 101 |
| SRR445726 (H. sapiens, RNA-Seq) | — | 49.2 | — | 100 | ERR532393 (metagen.) | — | 35.8 | — | 100 |
| SRR490961 (H. sapiens, RNA-Seq) | — | 49.1 | — | 100 | SRR065390 (C. elegans) | 103 | 67.6 | 66 | 100 |
| SRR1294116 (H. sapiens, RNA-Seq) | — | 46.0 | — | 101 | SRR689233 (M. musc., RNA-seq) | — | 16.4 | — | 90 |
| SRR490976 (H. sapiens, RNA-Seq) | — | 33.3 | — | 100 | SRR635193 | — | 27.3 | — | 54 |
| SRR445718 (H. sapiens, RNA-Seq) | — | 32.9 | — | 100 | MiSeq | 4.6 | 5.7 | 373 | 151 |
|  |  |  |  |  | SRR554369 (P. aeruginosa) | 6 | 1.7 | 50 | 100 |

The left part of the table presents the datasets used in the SE mode experiments and the right part the datasets used in the PE mode experiments. Additionally, in the SE experiments halves of the listed PE datasets were used (e.g., ERP001775\_1). The dataset sizes are given in the following tables. The column name “gen. length” denotes the length of the genome of the corresponding species. The genome length and coverage are not given for RNA-seq or metagenomic experiments. The name “MiSeq” denotes the dataset MiSeq\_Ecoli\_MG1655\_110721\_PF.

Tables 2 and 3 present the results in the order non-preserving and order-preserving SE modes, respectively (SE and SE_ORD in short). The table rows are sorted from the largest to the smallest dataset. The same hardware setup and essentially the same datasets were used in our earlier work (Kowalski and Grabowski, 2020), where also the results of Minicom and FQSqueezer can be seen. In both experiments, PgRC 1.2 wins in compression ratio over SPRING in 14 out of 15 cases. In the SE mode, the new version of PgRC wins over SPRING by 14.7% in the average compression ratio, and loses about 0.5% compared to v1.1, mostly due to the 6.6% loss on SRR870667_1 (on the remaining datasets the results are almost the same, with slight advantages for one side or the other). Please note that all the given averages here are geometric means, e.g., for a collection of three datasets, if algorithm X is twice faster than algorithm Y on some dataset (i.e., X’s ctime is half the ctime of Y) and also twice slower than Y on another dataset, and they have equal ctimes on the third dataset, then their ctime measures are equal on average.

**Table 2.** Compression, time and memory usage in the order non-preserving regime on SE datasets

| dataset | size<br>[Gbs] | SPRING |  |  |  |  | PgRC 1.1 |  |  |  |  | PgRC 1.2 |  |  |  |  |
| --- | --- | --- | --- | --- | --- | --- | --- | --- | --- | --- | --- | --- | --- | --- | --- | --- |
|  |  | ratio | ctime | dtime | cmem | dmem | ratio | ctime | dtime | cmem | dmem | ratio | ctime | dtime | cmem | dmem |
| ERP001775_1 | 44.39 | 0.2283 | 3511.5 | 302.7 | 23.48 | 5.27 | 0.2190 | 16943.0 | 247.6 | 38.20 | 16.38 | 0.2175 | 3510.2 | 277.4 | 38.41 | 16.38 |
| ERR174310_1 | 20.97 | 0.4087 | 1869.3 | 123.8 | 11.27 | 4.82 | 0.4509 | 14611.0 | 127.6 | 21.36 | 12.24 | 0.4484 | 2458.3 | 133.4 | 21.40 | 12.25 |
| SRR870667_1 | 7.48 | 1.2924 | 750.9 | 81.9 | 7.41 | 3.12 | 0.7655 | 12700.0 | 78.8 | 19.47 | 12.58 | 0.8157 | 1612.8 | 85.9 | 12.94 | 12.55 |
| SRR445724 | 5.09 | 0.4360 | 434.4 | 24.7 | 3.53 | 2.47 | 0.3401 | 2388.0 | 27.0 | 7.10 | 2.56 | 0.3432 | 477.5 | 28.3 | 2.72 | 2.56 |
| SRR445726 | 4.92 | 0.3880 | 440.1 | 21.7 | 3.60 | 2.49 | 0.3090 | 2071.9 | 24.3 | 6.39 | 2.24 | 0.3114 | 423.5 | 25.3 | 2.59 | 2.21 |
| SRR490961 | 4.91 | 0.2183 | 369.2 | 16.5 | 2.94 | 2.15 | 0.1794 | 1147.7 | 16.4 | 2.79 | 1.09 | 0.1793 | 278.4 | 16.9 | 2.49 | 1.11 |
| SRR1294116 | 4.64 | 0.2763 | 207.7 | 19.2 | 3.16 | 2.35 | 0.2333 | 1320.1 | 17.0 | 3.82 | 1.26 | 0.2329 | 254.6 | 17.9 | 2.42 | 1.25 |
| ERR532393_1 | 3.58 | 0.4325 | 178.3 | 16.6 | 3.11 | 1.85 | 0.3711 | 1582.8 | 18.6 | 5.25 | 1.27 | 0.3792 | 234.0 | 19.9 | 1.90 | 1.27 |
| SRR065390_1 | 3.38 | 0.2275 | 181.7 | 12.2 | 2.63 | 1.24 | 0.1858 | 720.3 | 10.8 | 1.89 | 0.76 | 0.1852 | 178.1 | 11.2 | 1.82 | 0.77 |
| SRR490976 | 3.33 | 0.3854 | 364.4 | 15.2 | 2.89 | 2.14 | 0.3316 | 1423.9 | 17.1 | 4.37 | 1.58 | 0.3306 | 326.5 | 18.0 | 1.80 | 1.57 |
| SRR445718 | 3.29 | 0.3371 | 225.7 | 14.6 | 2.63 | 2.03 | 0.2795 | 1168.9 | 15.0 | 3.35 | 1.15 | 0.2802 | 243.2 | 15.6 | 1.76 | 1.16 |
| SRR689233_1 | 1.48 | 0.1934 | 70.0 | 6.3 | 1.61 | 0.80 | 0.1687 | 288.8 | 4.7 | 0.76 | 0.31 | 0.1691 | 70.4 | 5.0 | 0.81 | 0.31 |
| SRR635193_1 | 1.47 | 0.2672 | 84.2 | 7.4 | 1.28 | 0.80 | 0.2558 | 381.0 | 5.7 | 1.66 | 0.49 | 0.2558 | 77.5 | 6.1 | 0.98 | 0.49 |
| MiSeq_1 | 0.87 | 0.1253 | 27.1 | 3.3 | 1.65 | 0.84 | 0.0937 | 102.3 | 1.9 | 0.36 | 0.09 | 0.0937 | 30.3 | 2.0 | 0.38 | 0.18 |
| SRR554369_1 | 0.17 | 0.2397 | 6.2 | 0.9 | 0.86 | 0.32 | 0.2393 | 29.4 | 0.6 | 0.20 | 0.05 | 0.2383 | 7.3 | 0.7 | 0.22 | 0.16 |

Compression ratios are in bits per base (bpb). “ctime” (resp. “dtime”) stands for compression (resp. decompression) time, and is expressed in seconds. “cmem” (resp. “dmem”) stands for peak compression (resp. decompression) memory usage and is expressed in gigabytes ( $\approx 10^9$ bytes).

**Table 3.** Compression, time and memory usage in the order preserving regime on SE datasets

| dataset | size<br>[Gbs] | SPRING |  |  |  |  | PgRC 1.1 |  |  |  |  | PgRC 1.2 |  |  |  |  |
| --- | --- | --- | --- | --- | --- | --- | --- | --- | --- | --- | --- | --- | --- | --- | --- | --- |
|  |  | ratio | ctime | dtime | cmem | dmem | ratio | ctime | dtime | cmem | dmem | ratio | ctime | dtime | cmem | dmem |
| ERP001775_1 | 44.39 | 0.5275 | 3621.0 | 344.9 | 23.48 | 4.07 | 0.5052 | 21997.0 | 339.8 | 41.72 | 19.46 | 0.5031 | 7793.0 | 372.4 | 41.73 | 19.46 |
| ERR174310_1 | 20.97 | 0.6953 | 1599.8 | 142.9 | 11.27 | 4.16 | 0.7275 | 16721.0 | 189.4 | 24.85 | 13.69 | 0.7246 | 4216.0 | 193.4 | 24.87 | 13.70 |
| SRR870667_1 | 7.48 | 1.4547 | 649.6 | 83.0 | 6.90 | 2.31 | 1.0305 | 12728.0 | 97.3 | 20.03 | 12.79 | 1.0805 | 1627.0 | 104.5 | 13.49 | 12.75 |
| SRR445724 | 5.09 | 0.6568 | 389.2 | 28.6 | 2.90 | 1.12 | 0.5937 | 2466.7 | 38.0 | 7.51 | 2.72 | 0.5966 | 511.1 | 39.1 | 4.11 | 2.71 |
| SRR445726 | 4.92 | 0.6128 | 376.0 | 26.4 | 2.83 | 1.08 | 0.5601 | 2160.4 | 34.9 | 6.78 | 2.38 | 0.5624 | 452.7 | 35.6 | 3.90 | 2.35 |
| SRR490961 | 4.91 | 0.4570 | 253.1 | 22.0 | 2.83 | 0.83 | 0.4286 | 1218.0 | 26.6 | 3.35 | 1.24 | 0.4286 | 307.1 | 27.0 | 3.53 | 1.28 |
| SRR1294116 | 4.64 | 0.5122 | 142.2 | 22.1 | 2.70 | 0.90 | 0.4740 | 1371.5 | 26.1 | 4.19 | 1.40 | 0.4736 | 281.8 | 26.9 | 3.45 | 1.38 |
| ERR532393_1 | 3.58 | 0.6665 | 123.5 | 21.0 | 1.78 | 1.00 | 0.6155 | 1613.8 | 26.1 | 5.54 | 1.38 | 0.6235 | 243.3 | 27.3 | 2.89 | 1.38 |
| SRR065390_1 | 3.38 | 0.4776 | 134.4 | 15.3 | 2.63 | 0.83 | 0.4258 | 749.3 | 17.8 | 2.45 | 0.84 | 0.4251 | 187.0 | 18.1 | 2.57 | 0.84 |
| SRR490976 | 3.33 | 0.5979 | 310.6 | 18.9 | 2.23 | 1.11 | 0.5777 | 1471.6 | 24.1 | 4.64 | 1.68 | 0.5769 | 350.2 | 24.8 | 2.58 | 1.70 |
| SRR445718 | 3.29 | 0.5707 | 181.0 | 17.3 | 2.21 | 0.89 | 0.5282 | 1207.4 | 21.8 | 3.61 | 1.24 | 0.5290 | 263.1 | 22.3 | 2.50 | 1.24 |
| SRR689233_1 | 1.48 | 0.4488 | 47.2 | 7.0 | 1.61 | 0.74 | 0.4289 | 313.3 | 7.6 | 1.11 | 0.36 | 0.4292 | 79.3 | 7.8 | 1.17 | 0.36 |
| SRR635193_1 | 1.47 | 0.6964 | 64.0 | 9.8 | 1.14 | 0.62 | 0.6799 | 423.3 | 10.4 | 1.87 | 0.57 | 0.6800 | 99.9 | 10.8 | 1.94 | 0.56 |
| MiSeq_1 | 0.87 | 0.2711 | 18.6 | 3.1 | 1.65 | 0.83 | 0.2400 | 106.3 | 3.0 | 0.44 | 0.11 | 0.2400 | 32.4 | 3.0 | 0.45 | 0.20 |
| SRR554369_1 | 0.17 | 0.4413 | 5.0 | 1.0 | 0.86 | 0.32 | 0.4362 | 30.7 | 0.9 | 0.21 | 0.06 | 0.4352 | 7.6 | 1.0 | 0.22 | 0.17 |

Compression ratios are in bits per base (bpb). “ctime” (resp. “dtime”) stands for compression (resp. decompression) time, and is expressed in seconds. “cmem” (resp. “dmem”) stands for peak compression (resp. decompression) memory usage and is expressed in gigabytes ( $=10^9$ bytes).

**Table 4.** Compression, time and memory usage in the order non-preserving regime on PE datasets

| dataset | size<br>[Gbs] | SPRING |  |  |  |  | PgRC 1.1 |  |  |  |  | PgRC 1.2 |  |  |  |  |
| --- | --- | --- | --- | --- | --- | --- | --- | --- | --- | --- | --- | --- | --- | --- | --- | --- |
|  |  | ratio | ctime | dtime | cmem | dmem | ratio | ctime | dtime | cmem | dmem | ratio | ctime | dtime | cmem | dmem |
| ERP001775 | 88.78 | 0.2307 | 6538.0 | 640.9 | 46.62 | 6.22 | 0.2026 | 25147.0 | 911.6 | 46.80 | 24.69 | 0.2015 | 5906.0 | 676.9 | 46.83 | 24.67 |
| ERR174310 | 41.93 | 0.3177 | 3426.7 | 269.8 | 21.94 | 5.17 | 0.3062 | 18099.0 | 289.0 | 39.18 | 17.93 | 0.3043 | 3664.0 | 309.4 | 39.45 | 17.96 |
| ERR532393 | 7.15 | 0.5026 | 357.3 | 39.2 | 5.61 | 2.95 | 0.4106 | 2954.1 | 54.5 | 9.00 | 2.55 | 0.4171 | 466.1 | 56.9 | 3.98 | 2.55 |
| SRR065390 | 6.76 | 0.2679 | 376.2 | 25.5 | 3.78 | 1.84 | 0.2050 | 1312.7 | 30.1 | 3.39 | 1.84 | 0.2044 | 331.7 | 30.9 | 3.44 | 1.85 |
| SRR689233 | 2.95 | 0.2390 | 168.9 | 12.7 | 1.94 | 1.33 | 0.2388 | 582.1 | 15.9 | 1.46 | 0.87 | 0.2392 | 154.0 | 16.4 | 1.58 | 0.87 |
| SRR635193 | 2.94 | 0.3392 | 171.5 | 16.3 | 1.95 | 1.18 | 0.3697 | 733.1 | 20.9 | 2.66 | 1.33 | 0.3686 | 169.0 | 21.7 | 1.90 | 1.32 |
| MiSeq | 1.73 | 0.1995 | 61.8 | 6.7 | 1.86 | 1.50 | 0.1489 | 273.9 | 6.3 | 0.75 | 0.32 | 0.1488 | 75.5 | 6.4 | 0.79 | 0.36 |
| SRR554369 | 0.33 | 0.2221 | 11.5 | 1.7 | 0.94 | 0.57 | 0.2165 | 50.0 | 1.4 | 0.27 | 0.10 | 0.2158 | 13.7 | 1.6 | 0.29 | 0.19 |

Compression ratios are in bits per base (bpb). “ctime” (resp. “dtime”) stands for compression (resp. decompression) time, and is expressed in seconds. “cmem” (resp. “dmem”) stands for peak compression (resp. decompression) memory usage and is expressed in gigabytes ( $=10^9$ bytes).

**Table 5.** Compression, time and memory usage in the order preserving regime on PE datasets

| dataset | size<br>[Gbs] | SPRING |  |  |  |  | PgRC 1.1 |  |  |  |  | PgRC 1.2 |  |  |  |  |
| --- | --- | --- | --- | --- | --- | --- | --- | --- | --- | --- | --- | --- | --- | --- | --- | --- |
|  |  | ratio | ctime | dtime | cmem | dmem | ratio | ctime | dtime | cmem | dmem | ratio | ctime | dtime | cmem | dmem |
| ERP001775 | 88.78 | 0.3842 | 6712.0 | 687.8 | 46.62 | 4.44 | 0.3399 | 30906.0 | 860.7 | 50.31 | 26.07 | 0.3403 | 10711.0 | 783.2 | 50.35 | 26.08 |
| ERR174310 | 41.93 | 0.4633 | 3709.0 | 293.0 | 21.93 | 4.39 | 0.4388 | 20517.0 | 377.7 | 40.84 | 19.18 | 0.4367 | 5918.0 | 353.7 | 40.86 | 19.21 |
| ERR532393 | 7.15 | 0.6208 | 246.6 | 52.7 | 3.64 | 1.77 | 0.5235 | 2907.2 | 63.3 | 9.28 | 2.43 | 0.5300 | 467.3 | 59.3 | 4.41 | 2.56 |
| SRR065390 | 6.76 | 0.3933 | 266.3 | 28.6 | 3.78 | 1.50 | 0.3190 | 1300.0 | 43.5 | 3.60 | 1.27 | 0.3185 | 341.1 | 38.5 | 3.86 | 1.40 |
| SRR689233 | 2.95 | 0.3668 | 124.0 | 13.2 | 1.94 | 1.32 | 0.3538 | 580.2 | 19.7 | 1.73 | 0.65 | 0.3542 | 160.2 | 17.5 | 1.85 | 0.76 |
| SRR635193 | 2.94 | 0.5616 | 130.9 | 19.2 | 1.80 | 1.02 | 0.5382 | 753.4 | 27.6 | 2.88 | 0.94 | 0.5383 | 189.0 | 24.1 | 3.00 | 1.04 |
| MiSeq | 1.73 | 0.2738 | 43.7 | 6.7 | 1.86 | 1.48 | 0.2174 | 264.8 | 7.8 | 0.75 | 0.21 | 0.2173 | 76.6 | 7.3 | 0.79 | 0.33 |
| SRR554369 | 0.33 | 0.3222 | 8.6 | 2.0 | 0.94 | 0.57 | 0.3086 | 48.6 | 1.9 | 0.30 | 0.08 | 0.3079 | 14.2 | 1.8 | 0.30 | 0.20 |

Compression ratios are in bits per base (bpb). “ctime” (resp. “dtime”) stands for compression (resp. decompression) time, and is expressed in seconds. “cmem” (resp. “dmem”) stands for peak compression (resp. decompression) memory usage and is expressed in gigabytes ( $=10^9$ bytes).

Still focusing on the SE experiment, we observe that the new PgRC is from 3.4 to 7.9 (and 4.8 on average) times faster in compression than the old version, while in the decompression (which is single-threaded) it is by a few percent (6.5% on average) slower. SPRING is still faster than PgRC in the compression, but now the average gap is only 10%, with swings; on SRR490961 PgRC 1.2 is by one third faster while on SRR870667_1 it is over twice slower. Overall, PgRC is faster here on 6 and slower on 9 datasets. The speed of PgRC 1.2 and SPRING is also similar in the decompression, with 3.7% average win of PgRC. Memory-wise, PgRC is, on average, somewhat more frugal in the compression (by almost 28%), but unfortunately not on the largest datasets, and the loss on ERR174310_1 is close to two-fold. PgRC 1.2 is also more frugal than v1.1 (by over 30% on average), but not on the two largest datasets, where it needs slightly more RAM. The overall improvements in space usage are due to some low-level internal optimizations.

Switching to the SE_ORD mode, we can observe the same general trend, although now the results are somewhat less favorable for our compressor. PgRC 1.2 has 7.2% average compression gain over SPRING and 0.4% loss to PgRC 1.1. The speedup of the new PgRC in compression spans from a factor of 2.8 to 7.8 (4.4 on average), while it is slower in the decompression by 3.8%. It is also slower in compression (resp. decompression) than SPRING by about 64% (resp. 20%) on average. The new version usually spent less working space (by about 19% on average) than v1.1 in the compression, but not in the decompression when the average loss is over 11%. SPRING remains the space leader, which is especially striking for the largest dataset (23.5 GB and 4.1 GB of required space for compression/decompression, compared to 41.7 GB and 19.5 GB from PgRC 1.2). The average wins of SPRING are more moderate though, around 10% and 41%, respectively.

The last two experiments, with PE and PE_ORD modes, generally follow the pattern. PgRC 1.2 dominates in the compression ratio over SPRING on 6 (resp. 8) out of 8 datasets, with the average gain slightly above 10%. Its compression performance with respect to the previous version is practically unchanged here (up to 1.6% differences). In compression speed SPRING wins by 5% (PE) and 55% (PE_ORD) while in the decompression the average gaps are around 16%. PgRC 1.2 is about 4 times faster in compression than its previous version, and also by a few percent faster in the decompression in the PE_ORD mode, with 0.7% average loss in the PE mode. In memory usage we again can see that SPRING is usually more succinct on the largest datasets (especially in the decompression), but not necessarily so on smaller ones.

To sum up, we can say that PgRC 1.2 is faster than PgRC 1.1 by a factor of about 4–5 (using 12 threads on a 6-core/12-thread machine) in compression, and more or less preserves its performance according to the other measures (compression ratio, decompression speed, memory usage in the compression and decompression). SPRING is usually somewhat faster, especially the order-preserving modes, and uses less working space (especially in the decompression), but cannot quite hold a candle to PgRC in the compression ratio, with roughly 7–15% compression ratio differences.

Fig. 2 presents the fraction of total PgRC’s processing time per phase, using 1 and 12 threads, respectively. The total compression time for the presented dataset, SRR065390, in the PE_ORD mode, is 1066.8s (resp. 341.1s) for 1 thread (resp. 12 threads). Using 12 threads gives thus a speedup by a factor of 3.1 on this dataset. We point out that the relative impact of reading the input data (from a HDD) grew significantly: more than 3 times. The read set division is over 5 times faster with 12 threads, which translates to a reduction from 40.3% to 24.6% for the fraction of the total time. Building the pseudogenomes using 12 threads is faster over 3.3 times, but the LZMA compression on them only twice (which is not bad, considering only 2 threads that LZMA uses). However, the pg compression time for PgRC 1.2 even with 1 thread is almost 10 times shorter than the corresponding time for PgRC 1.1. This spectacular improvement was possible due to the variable-length encoding phase, which is almost free in the processing time, and not only shortens the input for the LZMA, but also speeds up match searching over it, thanks to its more “dense” representation. To give another perspective, we note that in v1.1 the time fraction for this stage, i.e., “pg sequence, compression time” (cf. Fig. 3 in the Supplementary Material to (Kowalski and Grabowski, 2020)), was 23.5%; while in v1.2 it drops to 3.0% (1 thread) and 4.5% (12 threads), respectively. Finally, preparing the read alteration data is by a factor of 5.8 faster with 12 threads (the compression time is however unchanged). The relative impact of this stage, with regard to the total compression time, dropped from 22.9% (1 thread) to 13.7% (12 threads).

**Fig. 2.**
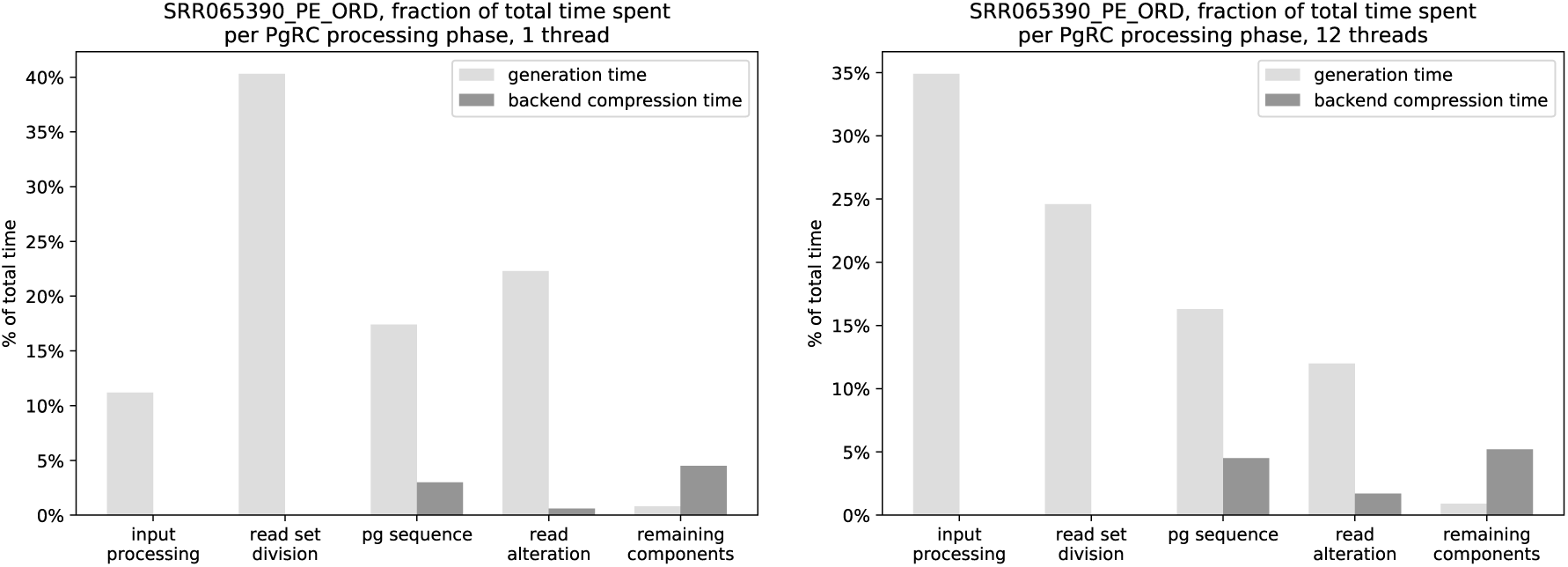
Fraction of the total time of the PgRC 1.2 compression process per phase, for SRR065390_PE_ORD (i.e., compressing SRR065390 in the order preserving PE mode). The left (resp. right) figure presents results for 1 thread (resp. 12 threads). The bars are doubled: the left of each pair concerns producing “raw” components, and the corresponding right one compressing it with LZMA or PPMd. The phase “pg compression” includes also the variable-length encoding, which takes about 0.5s for our data. The “read alteration” is an umbrella term to comprise operations producing and compacting alignment data: RC vs direct alignment flags, mismatch counts per read, mismatch symbols and mismatch positions within a read. The term “remaining components” concerns raw read positions (ordered) and read pairing data (for the PE_ORD mode).

**Fig. 3.**
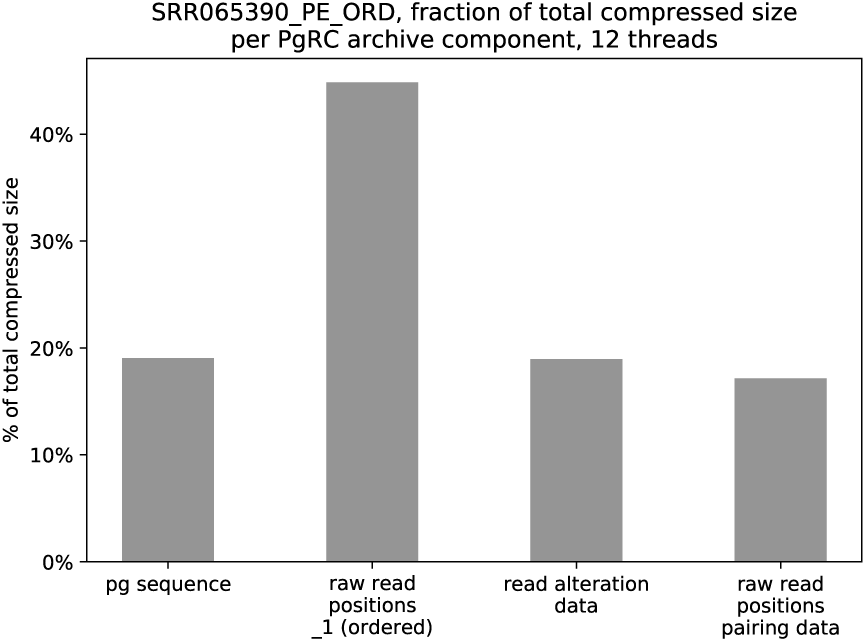
Fraction of the total size of the PgRC 1.2 compression process per component, for SRR065390_PE_ORD (i.e., compressing SRR065390 in the order preserving PE mode). The “read alteration data” is an umbrella term to comprise alignment data: RC vs direct alignment flags, mismatch counts per read, mismatch symbols and mismatch positions within a read.

Fig. 3 presents the fraction of total PgRC’s archive size per component. 12 threads were used here. Those results are almost identical as for PgRC 1.1 (cf. Fig. 4 in the Supplementary Material to (Kowalski and Grabowski, 2020)). Note that the raw positions of the reads from the first file (_1), which are stored in the input FASTQ order, use 2.61 times more space (the second bar) than the extra information required to decode the _2 file pairing (the fourth bar). There is an equilibrium between the compressed pseudogenomes’ size and the compressed sizes of the mapping data and the reverse-complement match data. It may be possible to make the sum of the (compressed) lengths of *P G*_*hq*_, *P G*_*lq*_ and *P G*_*N*_ smaller, with allowing more mismatches between *P G*_*hq*_ and mapping reads (and maybe also stricter quality requirements for the reads forming *P G*_*hq*_), but then the “read alteration data” grow, with the net result likely being a compression loss.

## 4 Conclusions

PgRC 1.2 is now a multi-threaded tool for compressing the DNA stream, improving the compression time of its previous version (v1.1) by a factor of about 4–5. One of the key ideas behind its success was to pack the DNA stream (from the pseudogenomes) using a variable-length encoding into bytes, which speeds up the later LZMA compression even more than could be expected from the more than threefold reduction in the length, thanks to making the data more ‘dense’ for the LZ77-like match finding process.

We note that PgRC is still work in progress, and some other engineering (and possibly algorithmic) improvements are possible. In particular, we are going to release a full-fledged FASTQ compressor, handling all data streams, including the quality scores and read headers.

